## Supplementary Materials, revised for "Evidence for increased parallel information transmission in human brain networks compared to macaques and mice"

* Corresponding authors

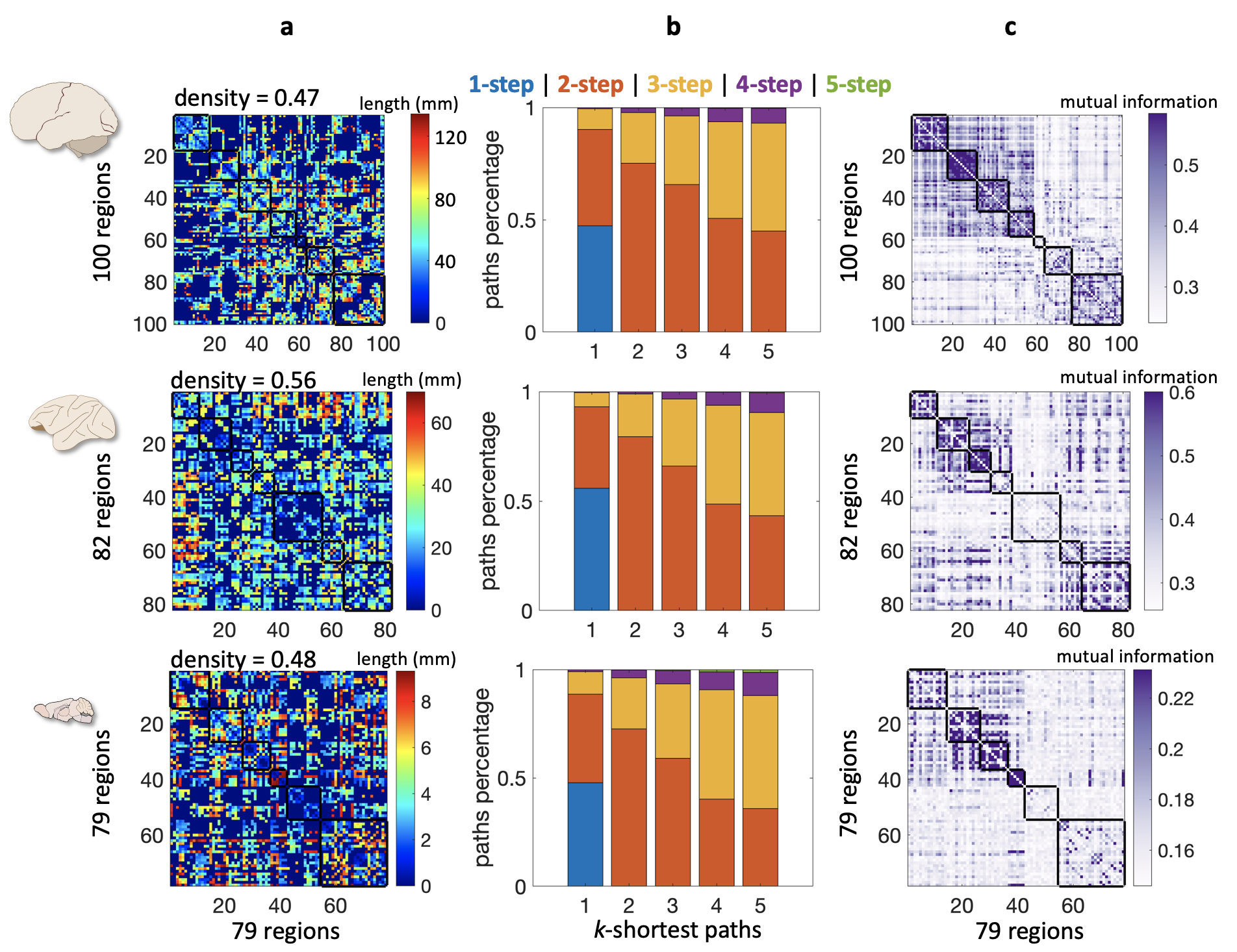

**Supplementary Figure 1. Cross-species brain structural and functional data.** Left: drawing of human, macaque, and mouse brains; each row in the figure corresponds to one species. (a) Group-representative structural connectivity matrices. The jet colormap represents the Euclidean distance between region pairs (in mm). Missing connections (i.e., disconnected region pairs) are represented in dark blue. The network density (i.e., the fraction of existing connections with respect to all possible connections) is reported above each matrix. (b) Percentage of *k*-shortest paths of length 1 (blue), 2 (orange), 3 (yellow), 4 (violet), or 5 (green) steps for the different matrices. Note the strong similarity between species. 1-step paths represent a minority of the overall *k*-shortest path ensembles in the three species (9%, 11% and 10%, respectively). (c) Group-average mutual information matrices (humans: n=100 biologically independent subjects; macaques: n=9 biologically independent subjects; mice: n=10 biologically independent subjects). For each species, brain regions are organised according to meaningful functional circuits (see Methods and Supplementary Figures 2, 3) which are highlighted by black squares along the matrices’ diagonals. Schematic of human, macaque, and mouse brains from scidraw.io (https://doi.org/10.5281/zenodo.3925945, https://doi.org/10.5281/zenodo.3926117, https://doi.org/10.5281/zenodo.3925909).

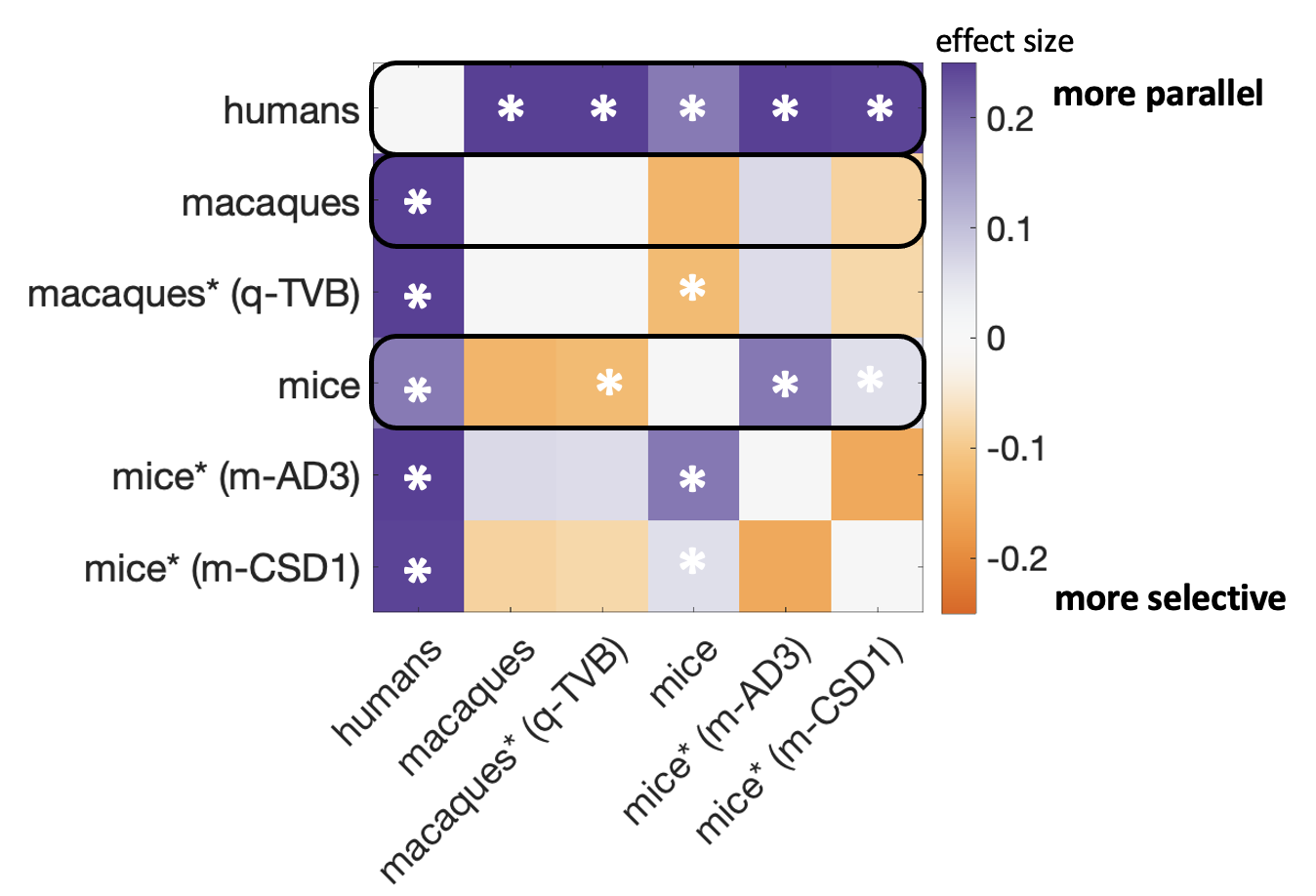

**Supplementary Figure 2. Parallel communication gap between humans and other mammalian species.** Results from pairwise statistical comparisons between group-average parallel communication scores (PCSs) of humans, macaques and mice (all main and replication datasets included). Main (non-anesthetized) and replication (anesthetized) datasets are reported on rows and columns (h-HCP: n=100 biologically independent subjects; q-NCS: n=9 biologically independent subjects; q-TVB: n=9 biologically independent subjects; m-GG: n=10 biologically independent subjects; m-AD3: n=10 biologically independent subjects; m-CSD1: n=51 biologically independent subjects); an asterisk next to the dataset name indicates that animals were anesthetized. The colorbar represents group-difference effect size, computed as difference of the medians normalized by the pooled median absolute deviation of the two groups: violet tones indicate that the row dataset has larger PCSs than the column dataset; orange tones indicate that the row dataset has smaller PCSs than the column dataset. White asterisks in the matrix indicate p-values < .05 / 15 (two-sided Mann-Whitney U tests, Bonferroni corrected).

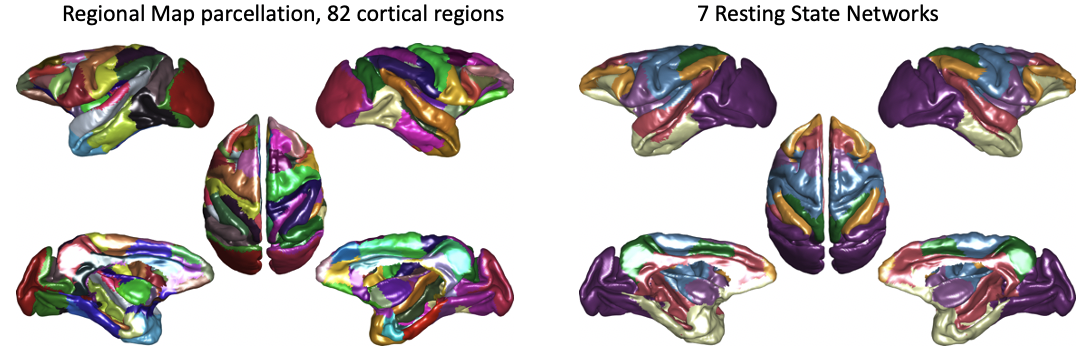

**Supplementary Figure 3. Assignment of macaque cortical brain regions to known functional systems** Each region of the Regional Map^1^ was assigned to one or more Brodmann areas, and then to one brain system (seven Yeo resting state networks^2^) using majority voting procedures. The cortical surface plots represent the 82 Regional Map regions (left; random color assignment) and their assignment to the seven brain systems (right; dark violet=visual; blue=somatomotor; green=dorsal attention; light violet=salience ventral attention; white=limbic; orange=executive control; red=default mode network). Brain networks were grouped into unimodal/multimodal systems (visual, somatomotor, dorsal attention, and salience ventral attention networks) and transmodal/limbic systems (executive control, default mode, and limbic networks)^3,4^.

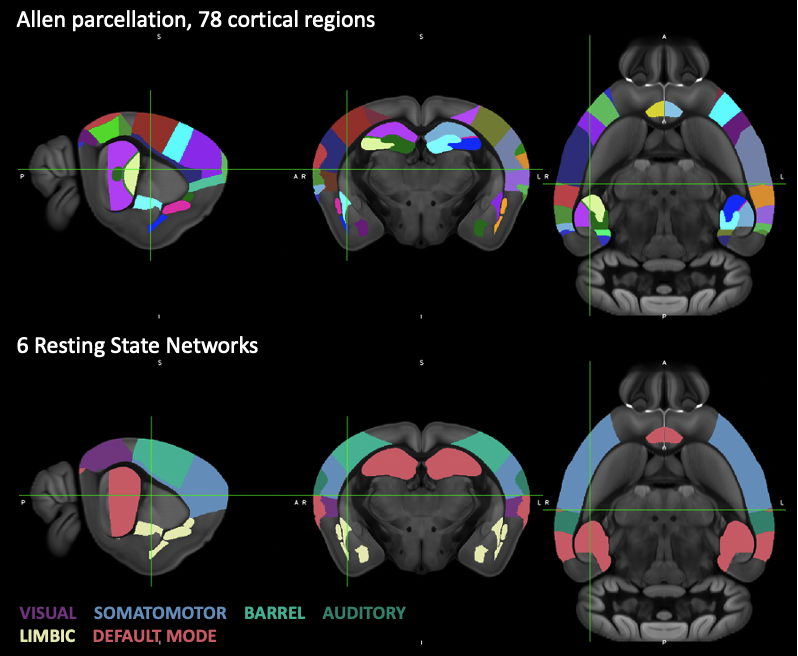

**Supplementary Figure 4. Assignment of mouse neocortical brain regions to known functional brain systems.** Each region of the Allen Mouse Brain Atlas (78 regions forming the isocortex, cortical subplate, and hippocampal formation were considered)^5^ was assigned to one out of six brain systems according to the functional mouse brain mapping of Zerbi and colleagues^6^. The Brain plots represent the 78 Allen Mouse Brain Atlas regions (top row; random color assignment) and their assignment to the six brain systems (bottom row; dark violet=visual; blue=somatomotor; light green=barrel; dark green=auditory; white=limbic; red=default mode network). Brain networks were grouped into unimodal systems (visual, somatomotor, barrel and auditory networks) and transmodal/limbic systems (default mode and limbic networks)^7^.

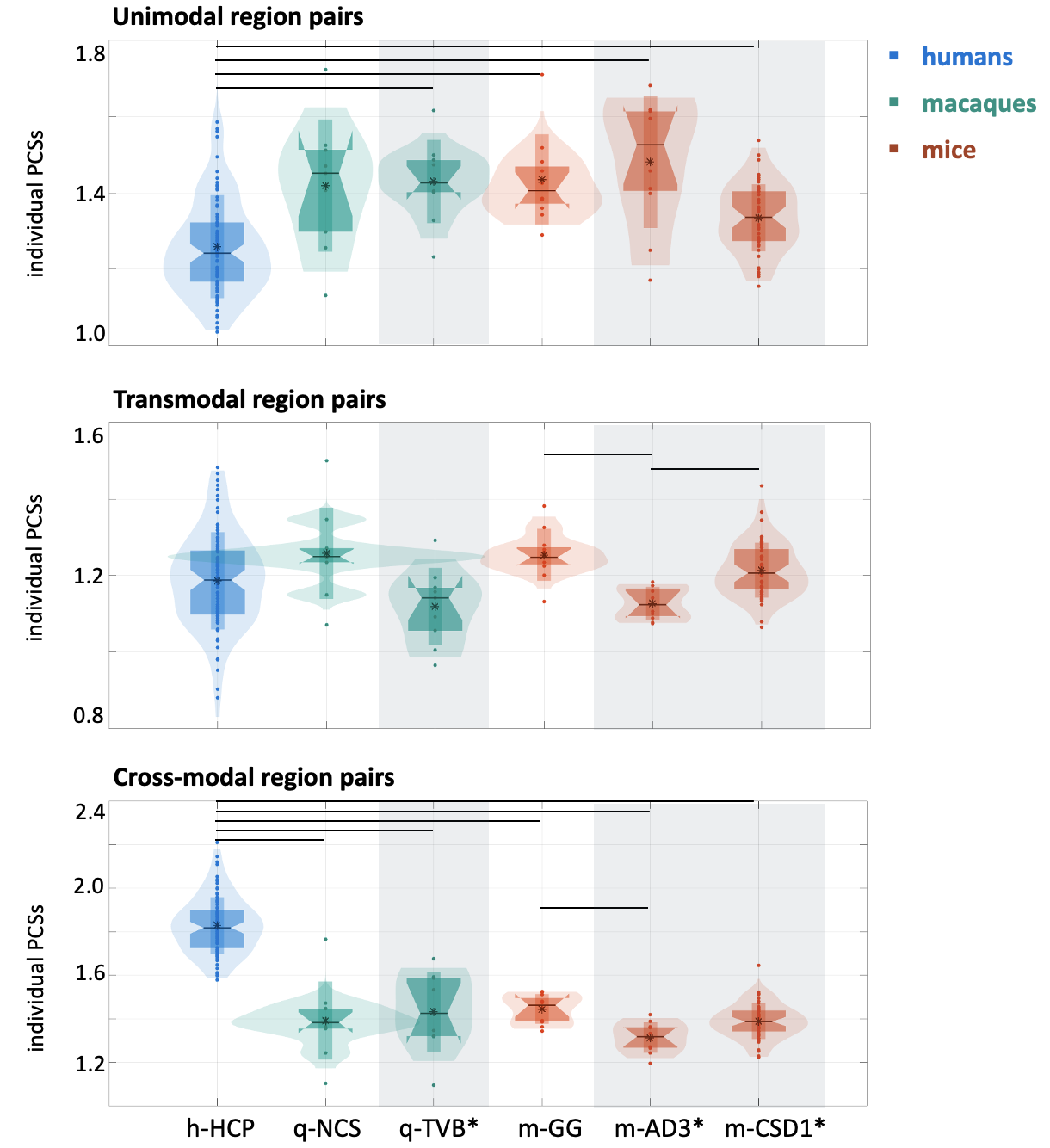

**Supplementary Figure 5. Individual parallel communication scores across species and brain systems.** Average parallel communication scores (PCSs) between unimodal systems (top row), between transmodal systems (middle row), and interconnecting unimodal and transmodal systems (‘cross-modal’ region pairs; bottom row) for individual experimental subjects of the three mammalian species (humans: blue; macaques: green; mice: brick red), and for all main and replication datasets (h-HCP: n=100 biologically independent subjects; q-NCS: n=9 biologically independent subjects; q-TVB: n=9 biologically independent subjects; m-GG: n=10 biologically independent subjects; m-AD3: n=10 biologically independent subjects; m-CSD1: n=51 biologically independent subjects). Statistically significant pairwise comparisons are indicated with horizontal black lines (two-sided Mann-Whitney U tests; p < .05/15, Bonferroni-corrected multiple comparisons). Datasets including anesthetized animals are indicated with a gray-shaded rectangle and an asterisk. In the box plots, each dot represents an individual; vertical bars indicate mean ± standard deviation; notch bars indicate median and 1^st^-3^rd^ quartiles; shaded areas indicate 1^st^-99^th^ percentiles.

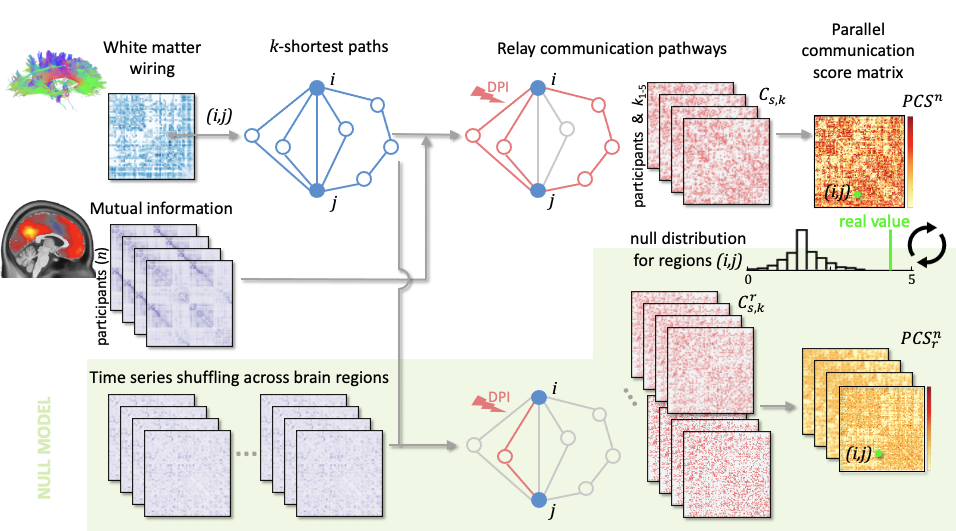

**Supplementary Figure 6. Methodological diagram detailing the assessment of parallel communication scores from real data and null model.** Top row: A weighted and symmetric structural connectivity matrix summarizes the white matter wiring of the brain for each species. For every pair of brain regions *(i, j)*, *k*=5 short structural paths (light blue) connecting the two regions are identified using the *k*-shortest path algorithm. Middle row: For every subject (human participant or animal), the mutual information between region pairs is computed from *z*-scored regional time courses obtained from fMRI recordings. By analyzing the mutual information values along each structural path, the data processing inequality (DPI) is used to assess whether the specific path represents an information-related pathway between regions *i* and *j* (light pink). For each subject *s*, *k* binary and symmetric communication matrices Cs,k are obtained, indicating whether the *k*-shortest path between a region pair is or is not a relay information-related pathway (light pink matrices). The sum of the *k* communication matrices gives an individual parallel communication score (PCS) matrix, with values between 0 and *k*. The group-average parallel communication matrix *PCS^n^* over *n* subjects is represented on the right (white-to-red colormap). Bottom row (green-shaded area): The null model is built by randomly shuffling the time courses of each subject over brain regions. The same above-described procedure is then applied to compute a group-average parallel communication score matrix for each randomization *r*. Note that the original time courses and group-average structural connectivity matrices are not modified by this null model. The randomization is repeated 3000 times for each subject to build a null distribution of group-representative parallel communication score matrices (represented on the right). Each entry of the original parallel communication score matrix *PCS^n^* (green line) is then compared to the corresponding null distribution by computing its *z*-score or its p-value with respect to the null distribution (black-and-white histogram). The p-values were computed by counting the percentage of surrogate PCS scores in the null distribution that exceeds the real PCS value.

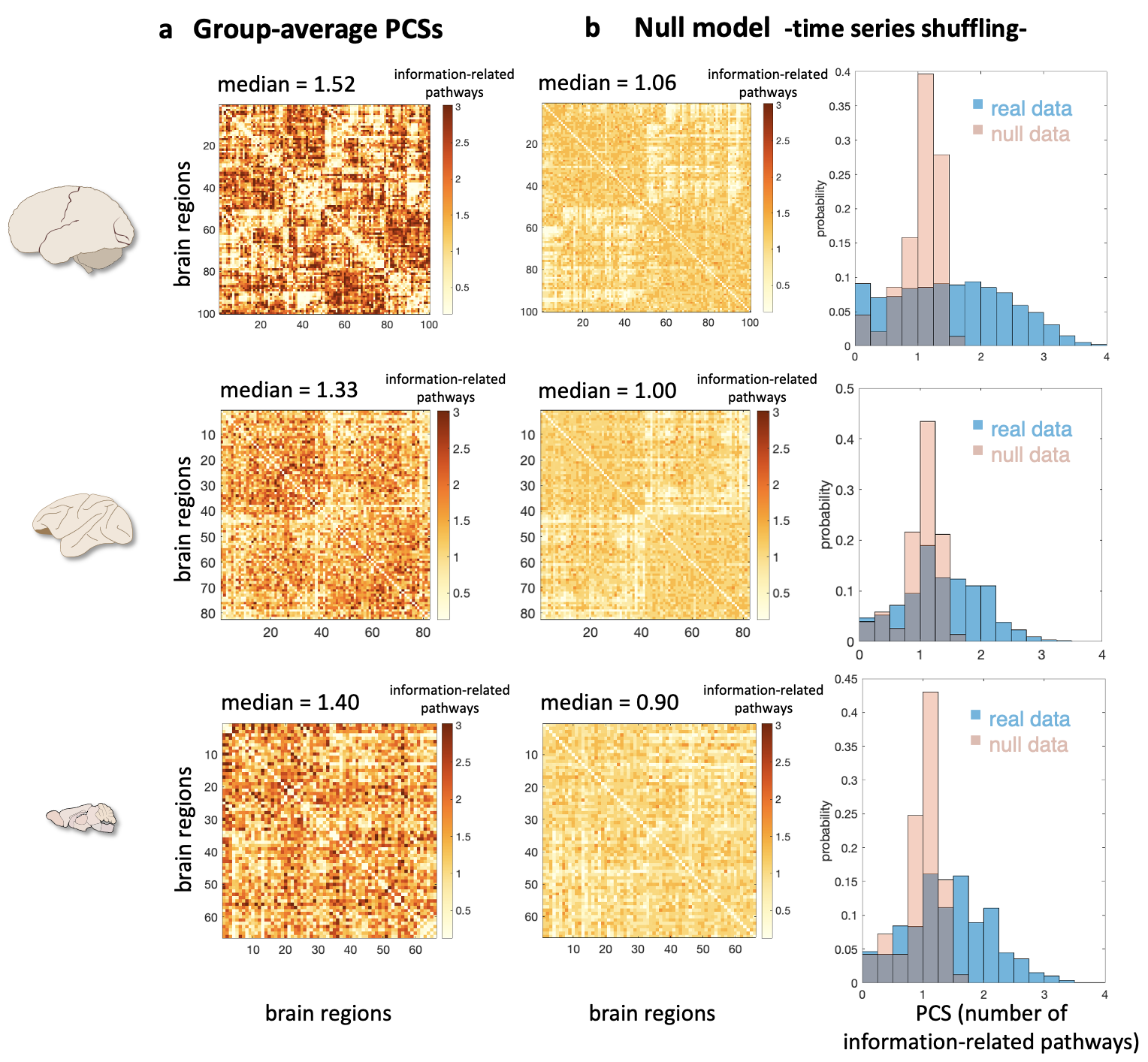

**Supplementary Figure 7. Real and null model group-average parallel communication scores.** Left: schematic of human, macaque, and mouse brains from scidraw.io (https://doi.org/10.5281/zenodo.3925945, https://doi.org/10.5281/zenodo.3926117, <https://doi.org/10.5281/zenodo.3925909>). Each row in the figure corresponds to one species. (a) Group-average parallel communication score (PCS) matrices representing PCSs between every pair of brain regions, averaged across individuals (similarly to Fig. 2; the color scale unit is the number of information-related pathways). (b) Null model: group-average parallel communication score matrices obtained by randomly shuffling the fMRI time series across brain regions and computing surrogate PCSs with unchanged structural connectome architecture (n=3000). The represented null-model matrices are the average over 3000 permutations. (c) Histograms of group-average PCS scores obtained from real data (blue) and null model (pink). For all the species, the real PCS histograms significantly differ from the null ones (two-sample Kolmogorov-Smirnov tests, p < .05), indicating that parallel communication levels are not the trivial by-product of structural connectivity architectures and multivariate statistical properties of fMRI time series. Humans: n=100 biologically independent subjects; macaques: n=9 biologically independent subjects; mice: n=10 biologically independent subjects

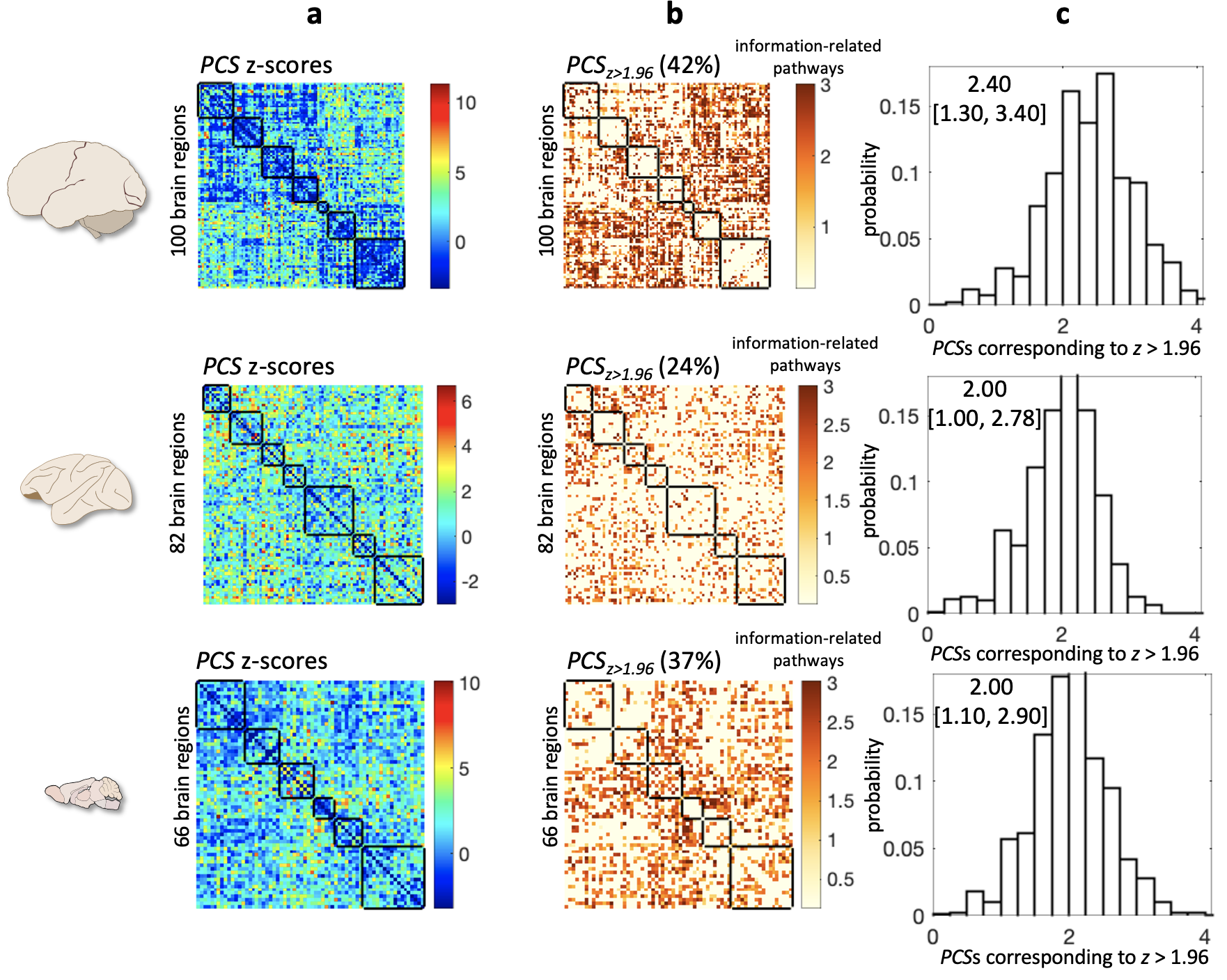

**Supplementary Figure 8. Significant parallel communication scores after z-score screening.** Left: schematic of human, macaque, and mouse brains from scidraw.io (https://doi.org/10.5281/zenodo.3925945, https://doi.org/10.5281/zenodo.3926117, <https://doi.org/10.5281/zenodo.3925909>). Each row in the figure corresponds to one species. (a) Matrices representing the z-scores of parallel communication scores (PCSs) with respect to the null distribution (n = 3000) obtained by randomly shuffling the fMRI time series across brain regions. (b) Significant PCSs values from z-score screening, i.e., selecting z-score values > 1.96. The percentage of region pairs with significant PCS is reported on top of each matrix (humans: 42%; macaques: 24%; mice: 37%). The color scale represents the PCS for significant region pairs. In (a) and (b), brain regions are organized according to meaningful functional circuits which are highlighted by black squares along the matrices’ diagonals (see Methods and Supplementary Figures 2, 3). (c) Histograms of significant group-average PCS scores (z-score > 1.96). Median [5-, 95-percentile] of significant PCS values for each species is reported atop each histogram. The cross-species distributions of significant PCSs were pairwise statistically different although the distance between macaque and mouse distributions was minor (two-sample Kolmogorov-Smirnov tests human-macaque: D_2087,794_ = 0.309, p < 10^-47^; human-mouse: D_2087,788_ = 0.297, p < 10^-43^; macaque-mouse: D_794,788_ = 0.099, p < 10^-3^), highlighting a gap between macaques and mice, with lower PCSs and mainly selective information transmission, and humans, with higher PCSs and presence of parallel communication. Only 10 human participants (instead of 100) were used for this analysis because the null distribution standard deviation (and therefore the z-scores) are affected by the number of participants, which we kept similar across species (n=10 biologically independent humans, n=9 biologically independent macaques, n=10 biologically independent mice).

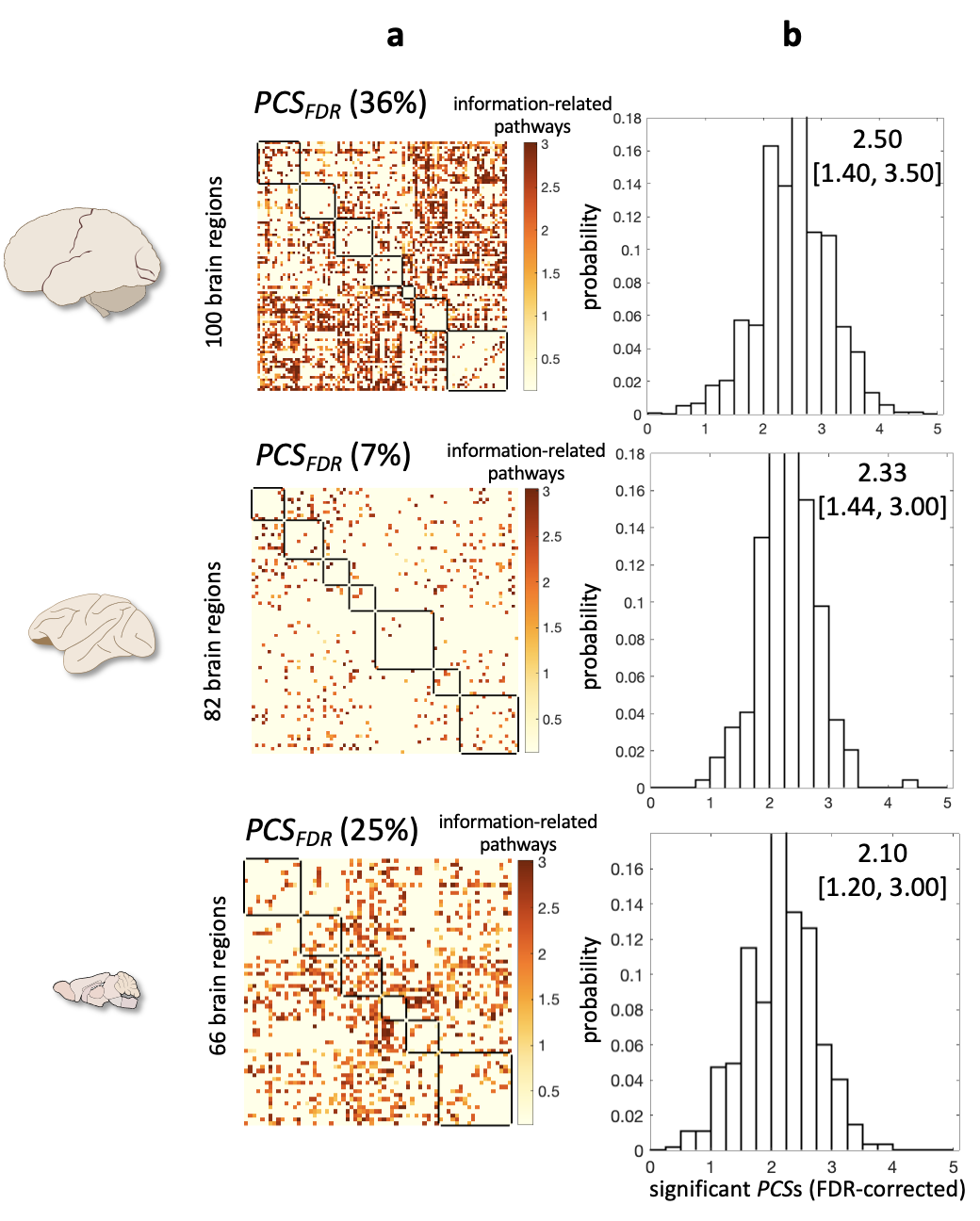

**Supplementary Figure 9. Significant parallel communication scores after FDR correction.** Left: schematic of human, macaque, and mouse brains from scidraw.io (https://doi.org/10.5281/zenodo.3925945, https://doi.org/10.5281/zenodo.3926117, https://doi.org/10.5281/zenodo.3925909). Each row in the figure corresponds to one species. (a) Significant PCSs values after false discovery rate (FDR) correction at FDR < .05. For each brain region pair, a p-value was computed by counting the percentage of surrogate PCS scores in the null distribution of that particular region pair that exceeds the real value. The color scale represents the PCS for significant region pairs. The percentage of region pairs with significant PCS (FDR-corrected p < .05) is reported on top of each matrix (humans: 36%; macaques: 7%; mice: 25%). Brain regions are organized according to meaningful functional circuits which are highlighted by black squares along the matrices’ diagonals (see Methods and Supplementary Figures 2, 3). (b) Histograms of significant group-average PCS scores (FDR-corrected p < .05). Median [5-, 95-percentile] of significant PCS values for each species is reported atop each histogram. The cross-species distributions of significant PCSs were pairwise statistically different (two-sample Kolmogorov-Smirnov tests human-macaque: D_1766,245_ = 0.238, p < 10^-10^; human-mouse: D_1766,547_ = 0.291, p < 10^-30^; macaque-mouse: D_245,547_ = 0.297, p < 10^-9^). Only 10 human participants (instead of 100) were used for this analysis because the null distribution standard deviation (and therefore the p-values) are affected by the number of participants, which we kept similar across species (n=10 biologically independent humans, n=9 biologically independent macaques, n=10 biologically independent mice).

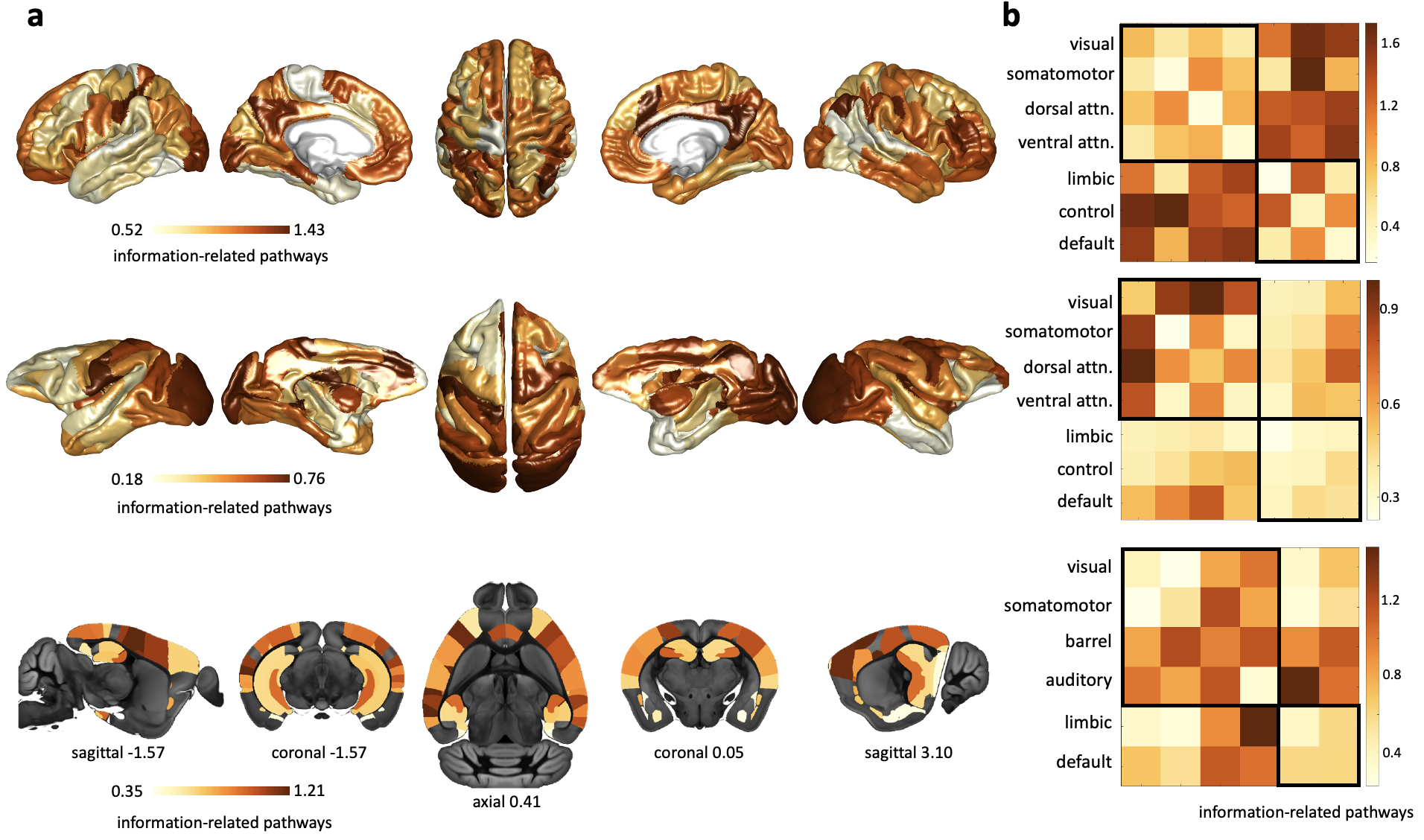

**Supplementary Figure 10. Brain topography of significant parallel communication scores after z-score screening.** (a) Cortical distributions of relay communication, quantified as the average significant parallel communication score (PCS) of each brain region with the rest of the brain network (first row: human, fsaverage6 cortical surface; second row: macaque, F99 template; third row: mouse, ABI template). Significant PCSs were identified by selecting z-score values > 1.96 compared to null distributions. For each species, the light yellow-to-brown colormap is scaled between the 5^th^ and 95^th^ percentiles of the cortical values, and it represents a nodal average number of information-related pathways. (b) Average significant PCSs within and between brain systems, for humans, macaques and mice. Brain systems have been organized into unimodal/multimodal regions (upper-left black square) and transmodal regions (lower-right black square). Only 10 human participants (instead of 100) were used for this analysis because the null distribution standard deviation (and therefore the z-scores) are affected by the number of experimental subjects, which we kept similar across species (n=10 biologically independent humans, n=9 biologically independent macaques, n=10 biologically independent mice).

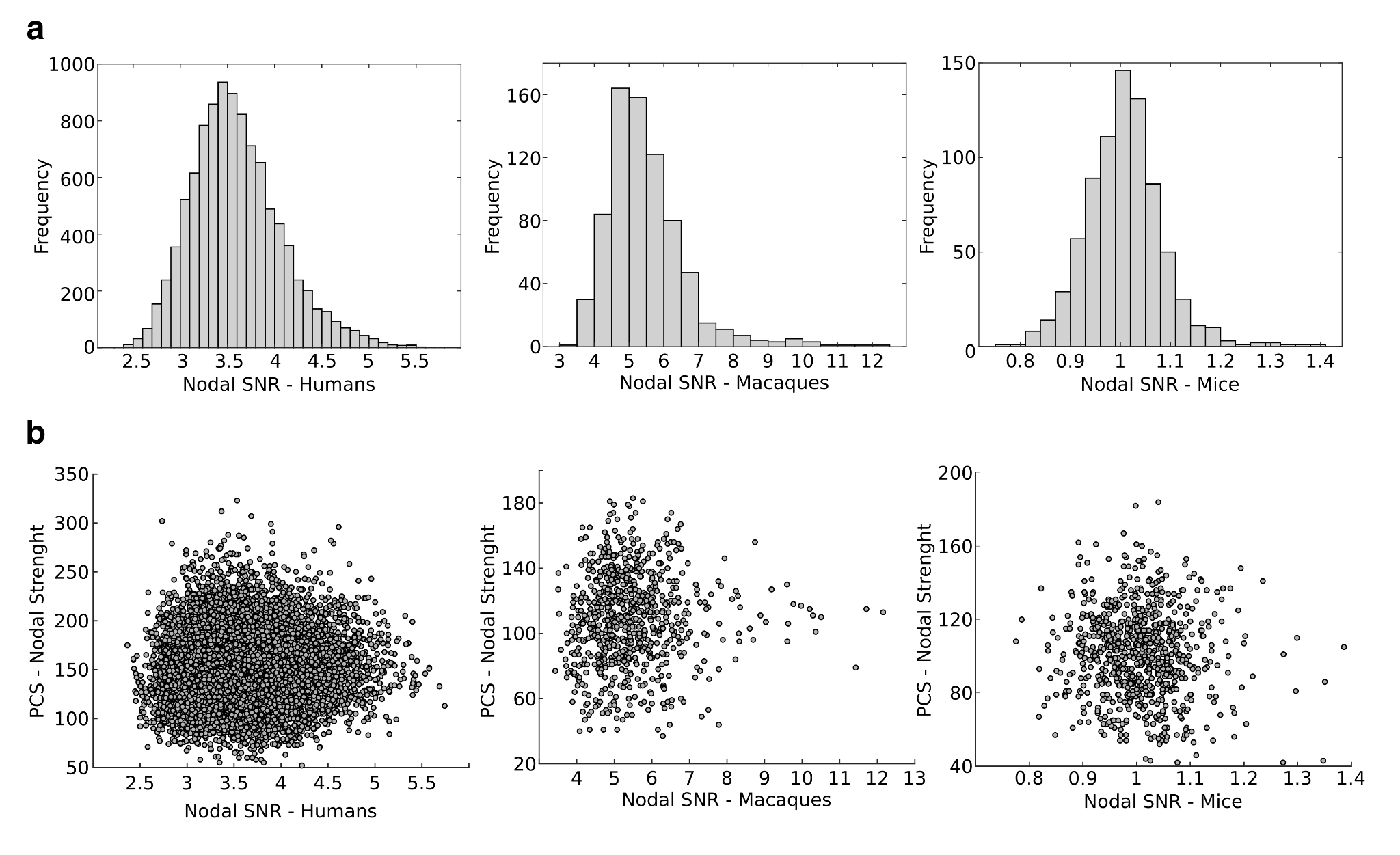

**Supplementary Figure 11. SNR evaluation with parallel communication scores across species.** (a) Nodal signal-to-noise ratio (SNR) distribution across species (humans: n=100 biologically independent subjects; macaques: n=9 biologically independent subjects; mice: n=10 biologically independent subjects). Due to the nature of the input data (i.e., parcellated nodal time series), SNR was defined as the ratio between low- and high-frequency power. (b) Scatter plots between individual nodal SNR and PCS nodal strength (i.e., sum over columns of the PCS matrices reported in Figure 2), for each species (h-HCP, q-TVB, m-AD3). Note how PCS does not correlate with SNR (specifically, in humans Spearman’s correlation coefficient ρ=0.0277; in macaques ρ=0.0782; in mice ρ=-0.0538). This finding demonstrates that basic scanning protocol differences between species can be excluded as a confounding factor for our brain communication results.

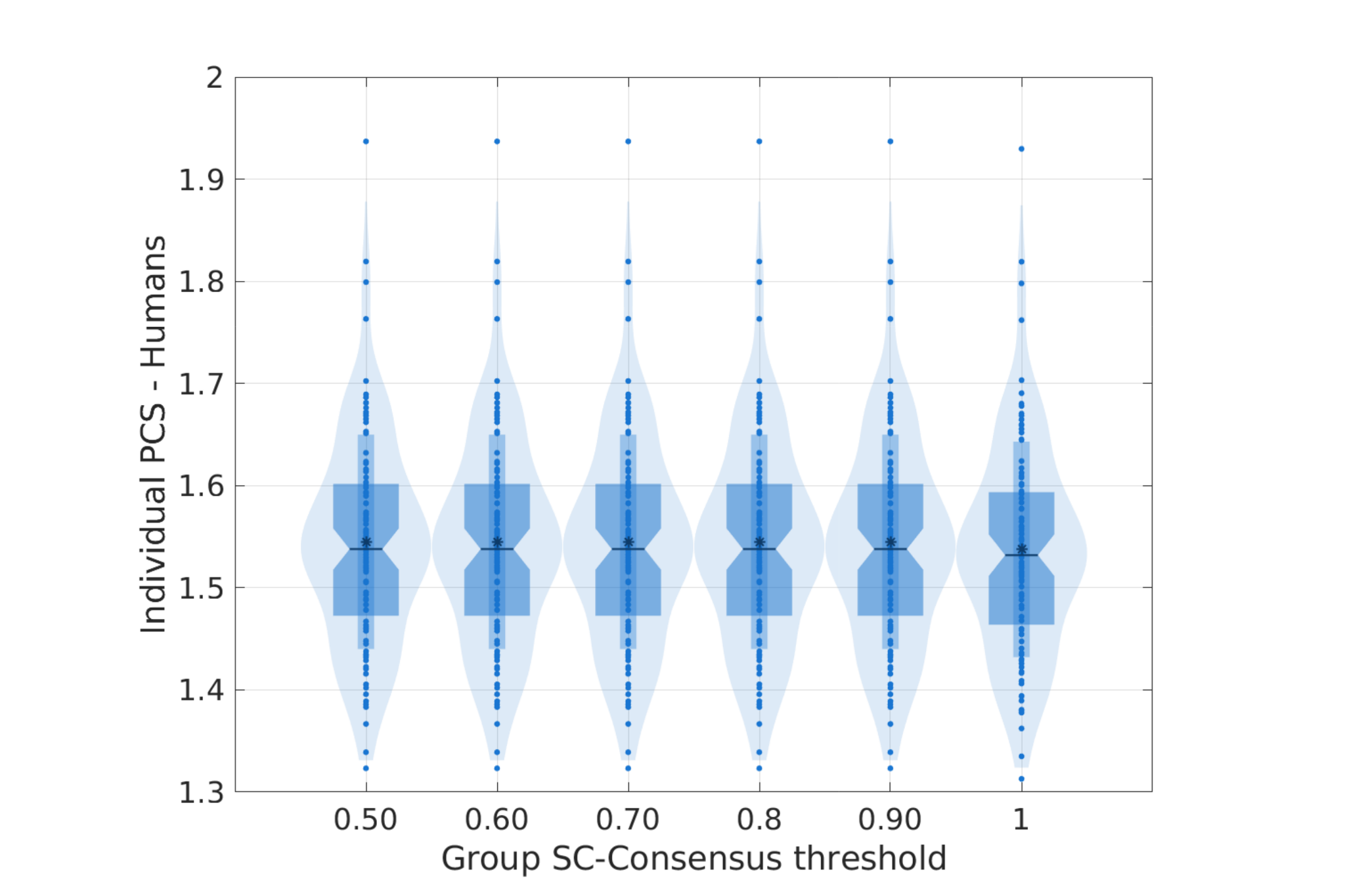

**Supplementary Figure 12. Relationship between parallel communication scores and structural network complexity.** The box plots depict individual parallel communication scores (PCSs) in humans (y-axis, h-HCP dataset, n=100 biologically independent subjects) when changing the group-representative structural connectome by tuning the consensus threshold from 100% (the value chosen for the main results) down to 50% (allowing a connection when present in half the cohort) in 10% increasing steps, thus progressively increasing the connectivity density (x-axis). In the box plots, each dot represents an individual; vertical bars indicate mean ± standard deviation; notch bars indicate median and 1^st^-3^rd^ quartiles; shaded areas indicate 1^st^-99^th^ percentiles. Note how individual PCSs are stable across the choices of the threshold, confirming that thresholding of the structural connectome and connectivity density -a simple indicator of network complexity- cannot explain the brain communication patterns found.

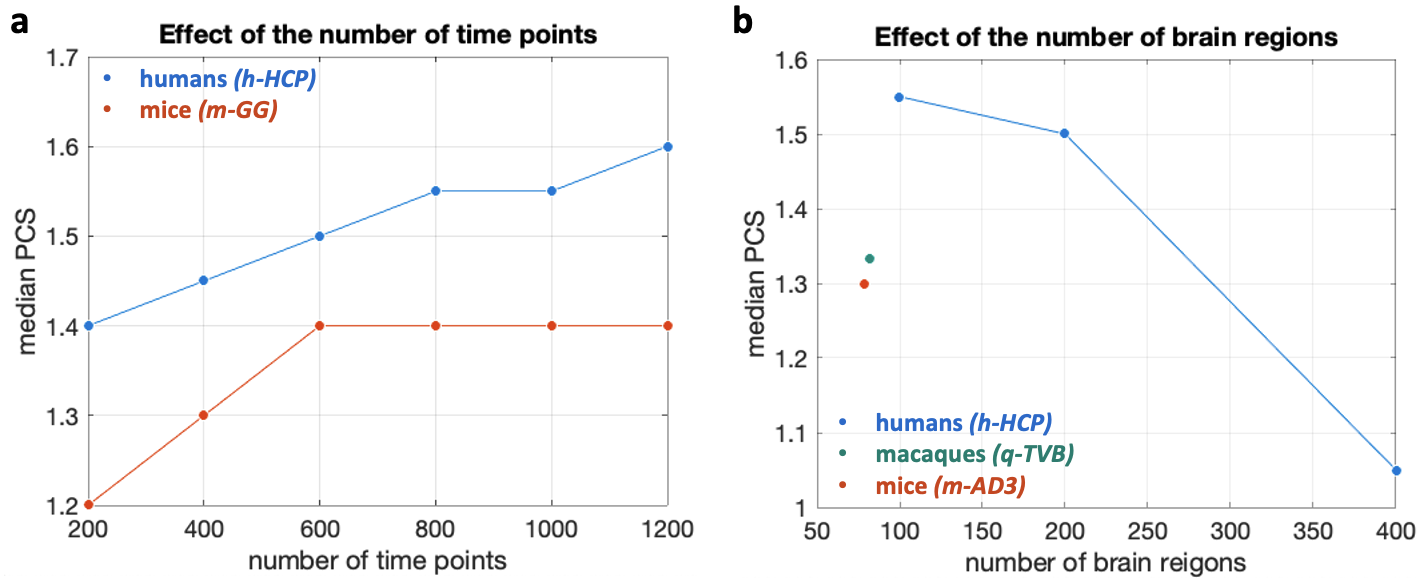

**Supplementary Figure 13. Sensitivity analysis: parallel communication scores and number of time points or brain regions.** (a) Each dot represents the median parallel communication score (PCS) computed from the group-average PCS matrices obtained from n=100 HCP human subjects (h-HCP dataset, light blue) and n=10 mice (m-GG dataset, brick red) as a function of the number of time points included in the analysis. We observe that PCS values increase with an increasing number of time points, reaching a plateau for longer scan durations. This analysis was limited to the datasets h-HCP and m-GG because of the availability of relatively long fMRI scans. (b) Each dot represents the median parallel communication score computed from group-average PCS matrices for the three species: n=100 HCP human subjects (h-HCP dataset, light blue), n=9 macaques (q-TVB dataset, green), and n=10 mice (m-AD3 dataset, brick red) as a function of the number of brain regions. We observe that, in humans, median PCS values tend to decrease with an increasing number of brain regions, although the variation is minor when comparing the cortical parcellations including 100 and 200 regions (Schaefer parcellations^8^).

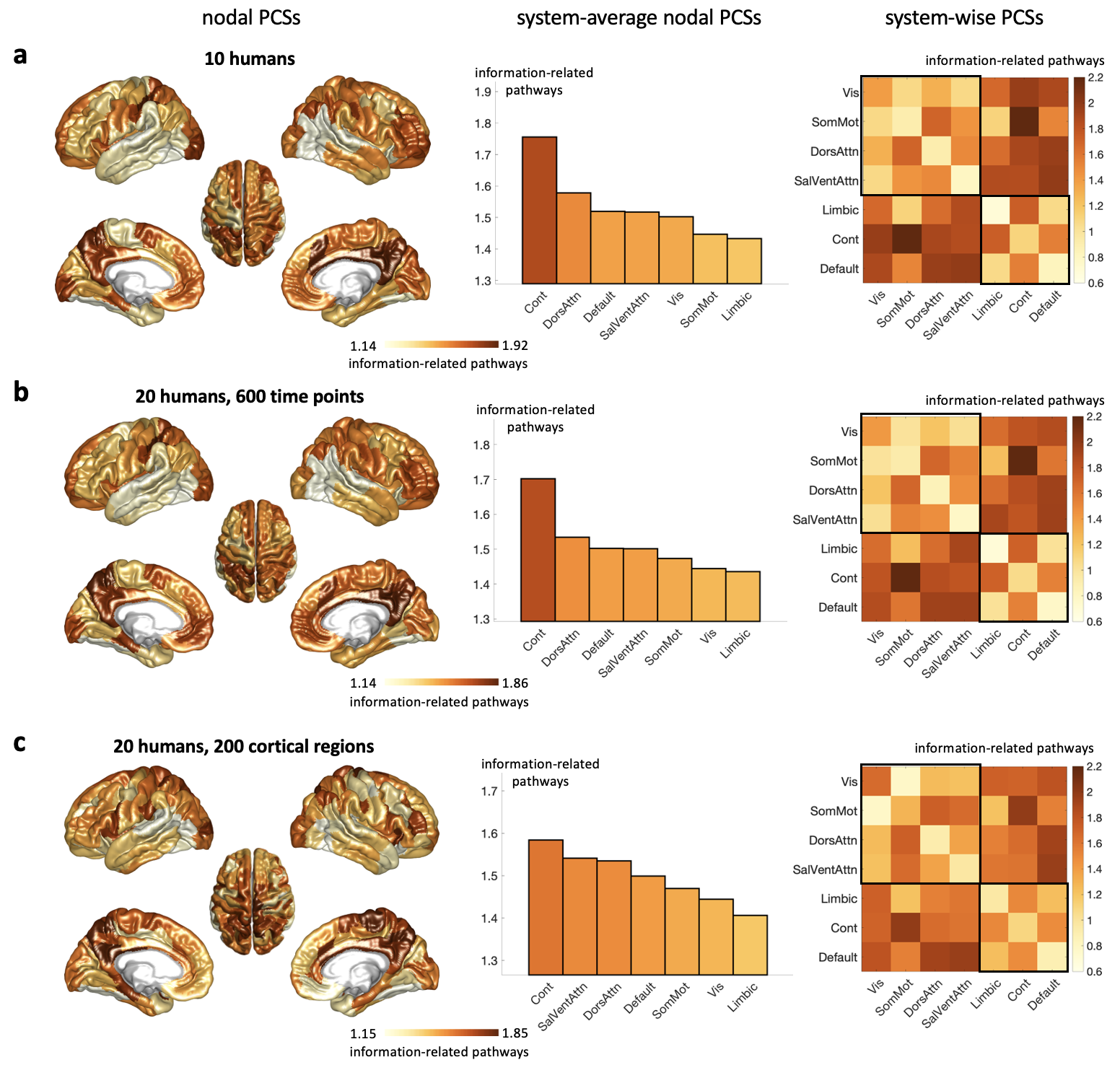

**Supplementary Figure 14. Sensitivity analysis: parallel communication topographies and number of time points or brain regions in humans.** Left: cortical distributions of relay communication, quantified as the average PCS of each brain region with the rest of the brain network (fsaverage6 cortical rendering). For each dataset, the light yellow-to-brown colormap is scaled between the 5^th^ and 95^th^ percentiles of the cortical values, with the color scale unit indicating the average number of information-related pathways. Middle: the average nodal communication scores per brain system are represented as bar plots. Right: average PCSs within and between brain systems, organized into unimodal/multimodal regions (upper-left black square) and transmodal regions (lower-right black square). HCP data: (a) n=10 biologically unrelated subjects, 100 cortical regions, 800 time points; (b) n=20 biologically unrelated subjects, 100 cortical regions, 600 time points; (c) n=20 biologically unrelated subjects, 200 cortical regions, 800 time points. Vis = visual; SomMot = somatomotor; DorsAttn = dorsal attention; SalVentAttn = salience ventral attention; Cont = executive-control; Default = default mode networks.

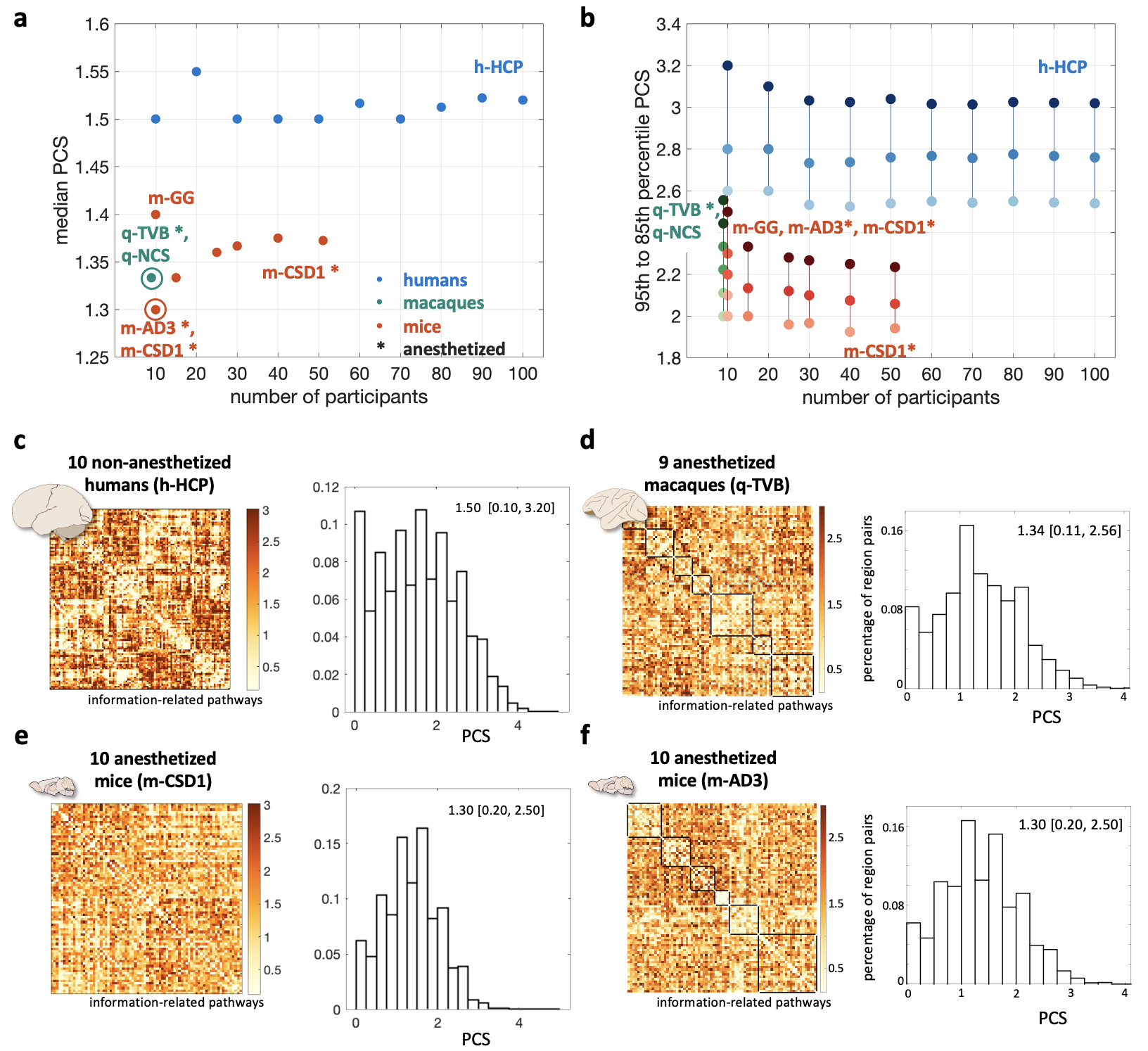

**Supplementary Figure 15. Sensitivity and replications analyses: group-average parallel communication scores and number of subjects.** The following datasets were included in these analyses: (i) [h-HCP] n=100 healthy adult human subjects from the HCP database^1^ - main dataset. Analyses were repeated while including different numbers of subjects (from 10 to 100 subjects); (ii) [q-TVB] n=9 anesthetized male adult macaque monkeys (8 Macaca mulatta, 1 Macaca fascicularis)^3^; (iii) [q-NCS] n=9 non-anesthetized male adult macaque monkeys (9 Macaca mulatta) from the PRIMatE Data Exchange initiative^4^ - main dataset; (iv) [m-AD3] n=10 anesthetized wild-type mice scanned at 6 months^5^; (v) [mCSD1] n=51 anesthetized wild-type mice scanned at 3 months^7^. Analyses were repeated while including different numbers of subjects (from 10 to 51); (vi) [m-GG] n=10 non-anesthetized wild-type adult mice scanned <6 months^9^ - main dataset. (a) Each dot represents the median parallel communication score (PCS) computed from the group-average PCS matrices obtained from the different datasets, as a function of the number of participants included in the analyses. Blue indicates human datasets, green macaque datasets, and brick red murine datasets; asterisks indicate datasets including anesthetized participants; a circle indicates superposition of PCS median values for multiple datasets (q-TVB and q-NCS; m-AD3 and m-CSD1 with 10 participants). (b) Largest PCS values for each dataset. Each set of three dots connected by a line represents the 95^th^, 90^th^ and 85^th^ percentiles of the PCS distribution obtained from the group-average PCS matrix of each dataset. (c-f) Group-average PCS matrices and their histograms for four representative datasets. The color scale unit for the PCS matrices is number of information-related pathways. (c) 10 human subjects (h-HCP); (d) 9 anesthetized macaques (q-TVB); (e) 10 anesthetized mice (m-CSD1); (f) 10 anesthetized mice (m-AD3). On the right, the histograms of the average PCS scores across region pairs highlight a gap between mice and macaques, with lower PCSs and mainly selective information transmission, and humans, with higher PCSs and presence of parallel communication. Median [5-, 95-percentile] PCS values for each species are reported atop each histogram.

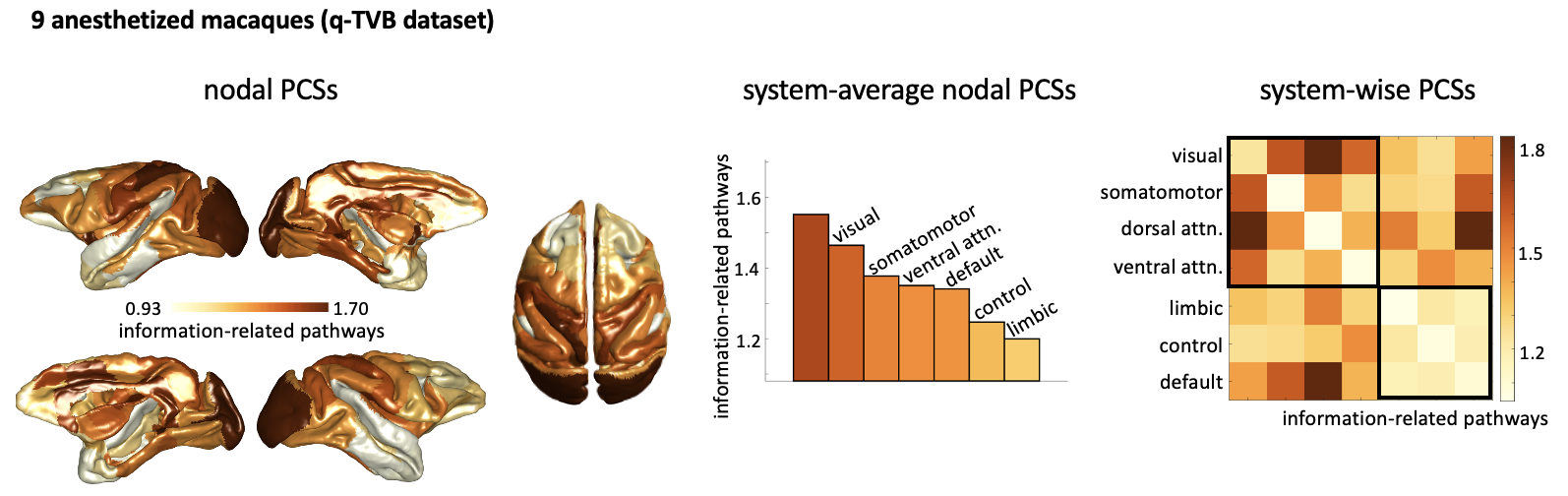

**Supplementary Figure 16. Replication analysis: parallel communication topography in macaques.** Data from n=9 anesthetized macaques (q-TVB dataset). Left: cortical distributions of relay communication, quantified as the average PCS of each brain region with the rest of the brain network (F99 template cortical rendering). The light yellow-to-brown colormap is scaled between the 5^th^ and 95^th^ percentiles of the cortical values, with the color scale unit indicating the average number of information-related pathways. Middle: the average nodal communication scores per brain system are represented as bar plots. Right: average PCSs within and between brain systems, organised into unimodal/multimodal regions (upper-left black square) and transmodal regions (lower-right black square).

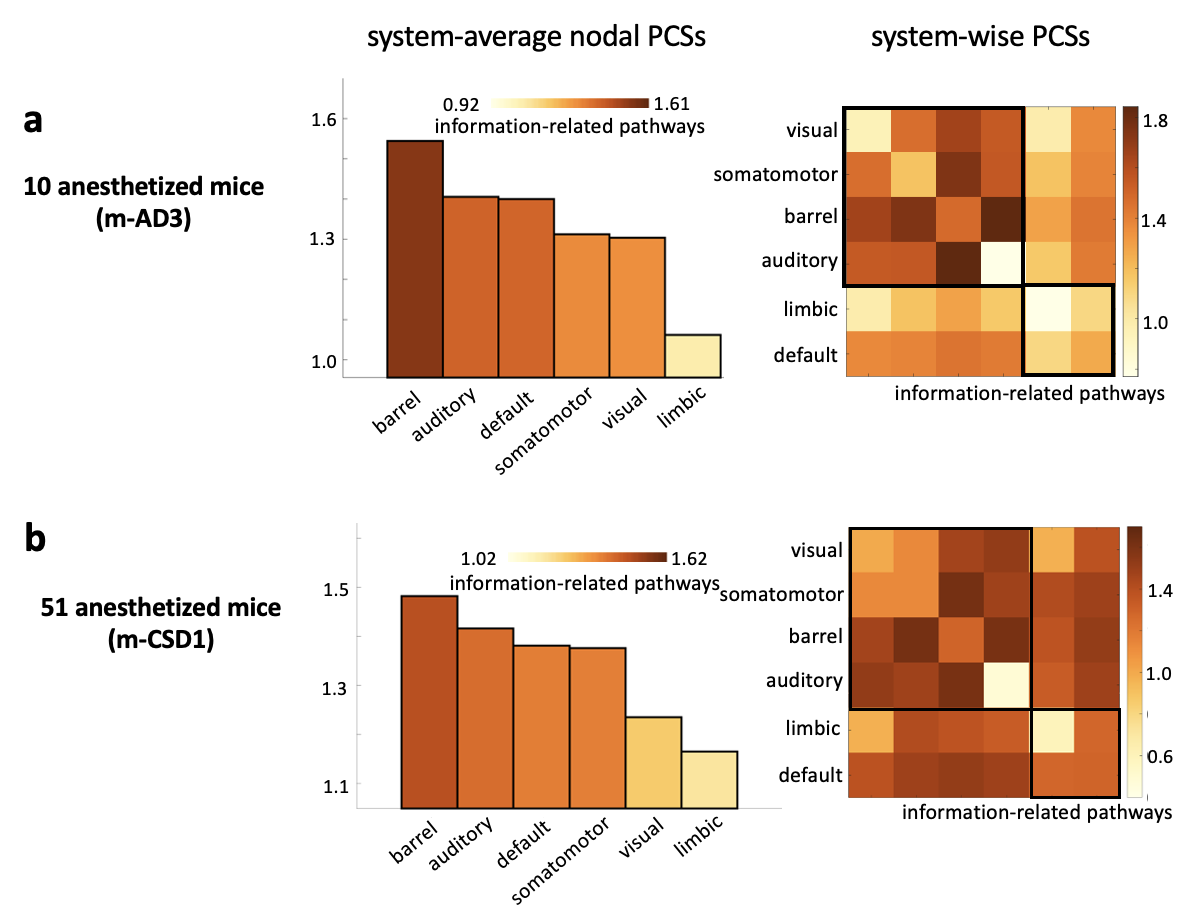

**Supplementary Figure 17. Replication analysis: parallel communication topography in anesthetized and awake mice.** Left: average nodal parallel communication scores (PCSs) per brain system. The light yellow-to-brown colormap is scaled between the 5^th^ and 95^th^ percentiles of the cortical values, with the color scale unit indicating the average number of information-related pathways. Right: average PCSs within and between brain systems, organized into unimodal regions (upper-left black square) and transmodal regions (lower-right black square). (a) n=10 anesthetized mice (m-AD3 dataset); (b) n=51 anesthetized mice (m-CSD1 dataset).

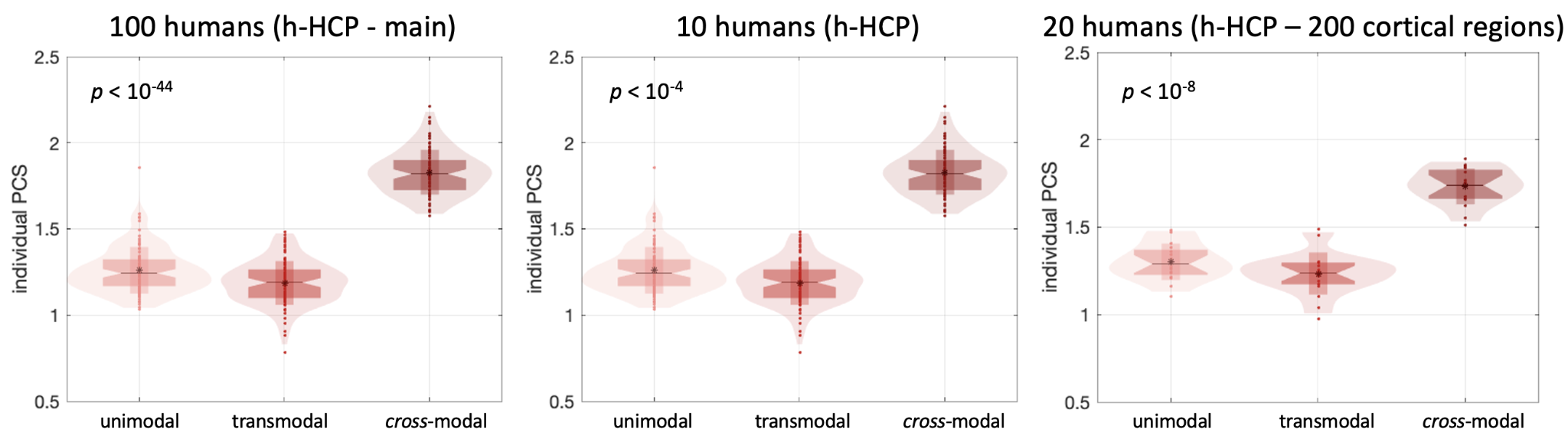

**Supplementary Figure 18. Sensitivity analysis: unimodal / transmodal / cross-modal communication and number of subjects or brain regions in humans.** Average parallel communication scores (PCSs) between unimodal systems, between transmodal systems, and interconnecting unimodal and transmodal systems (cross-modal) for individual human subjects. The following subject groups were considered: (left) human h-HCP main dataset (n=100 biologically unrelated subjects, 100 brain regions), (middle) n=10 biologically unrelated subjects, 100 brain regions, (right) n=20 biologically unrelated subjects, 200 brain regions. In the box plots, each dot represents an individual; vertical bars indicate mean ± standard deviation; notch bars indicate median and 1^st^-3^rd^ quartiles; shaded areas indicate 1^st^-99^th^ percentiles. Kruskal Wallis p-values for within-species comparisons are reported, testing the null hypothesis that unimodal, transmodal and cross-modal PCS scores originate from the same distribution.

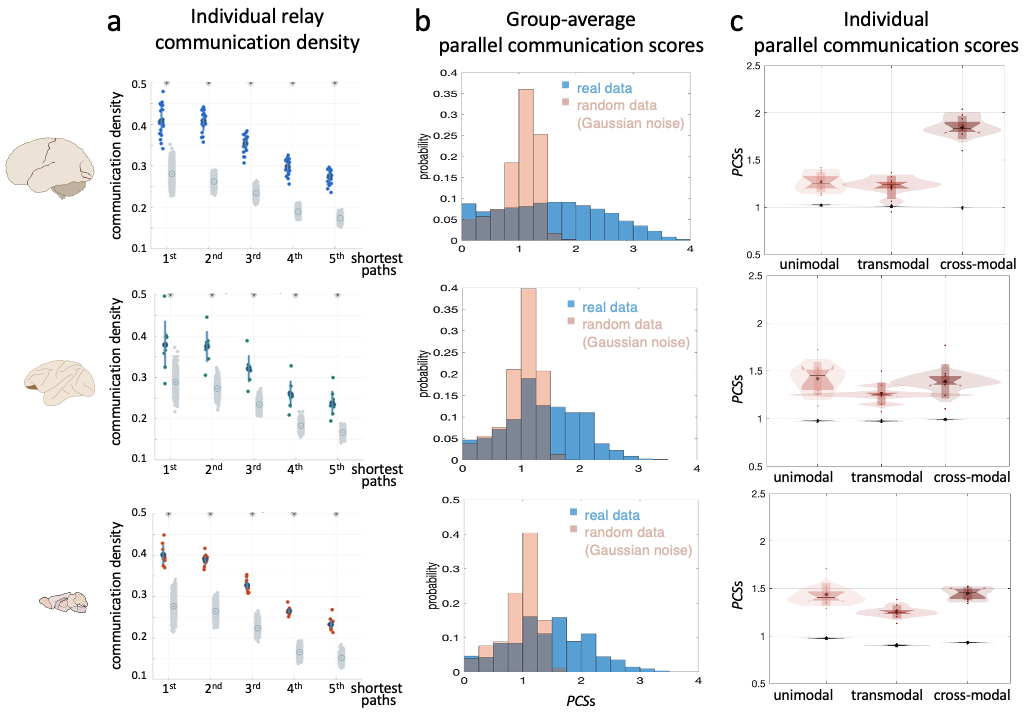

**Supplementary Figure 19. Comparison between real data and iid Gaussian noise null model.** Left: schematic of human, macaque, and mouse brains from scidraw.io (https://doi.org/10.5281/zenodo.3925945, https://doi.org/10.5281/zenodo.3926117, https://doi.org/10.5281/zenodo.3925909). Each row in the figure corresponds to one species. (a) Box plots representing the percentage of short structural paths in individual brain networks respecting the data processing inequality (communication density). Each coloured dot represents an individual. Grey dots represent null distributions obtained by plugging-in iid Gaussian noise at the nodes of the brain network; the brain network structural topology was unchanged (Methods). Circles and vertical bars indicate mean ± one standard deviation across individuals and randomizations. Paths are grouped according to the 1^st^ up to the 5^th^ shortest path between region pairs. (b) Histograms of group-average parallel communication scores (PCSs) obtained from real data (blue) and Gaussian null model (pink). For all the species, the real PCS histograms significantly differ from the null ones (two-sample Kolmogorov-Smirnov tests, p < .05), indicating that parallel communication levels are not the trivial byproduct of the structural network topology (weighted by the Euclidean distance between region pairs) and the brain spatial embedding preserved in the null model. Yet, the effect of the structural network topology is evident in the fact that the median PCSs in case of Gaussian noise are larger than zero. (c) Average PCSs between unimodal systems, between transmodal systems, and between unimodal and transmodal systems (cross-modal communication) for individual experimental subjects. In the box plots, each dot represents a subject individual; vertical bars indicate mean ± standard deviation; notch bars indicate median and 1^st^-3^rd^ quartiles; shaded areas indicate 1^st^-99^th^ percentiles. Values from the Gaussian null model are reported in the same panels as grey box plots. For all species and brain systems, the real individual PCS significantly differ from the null ones, indicating that the structural network topology cannot explain system-dependent spatial communication patterns. Humans: n=100 biologically independent subjects; macaques: n=9 biologically independent subjects; mice: n=10 biologically independent subjects.

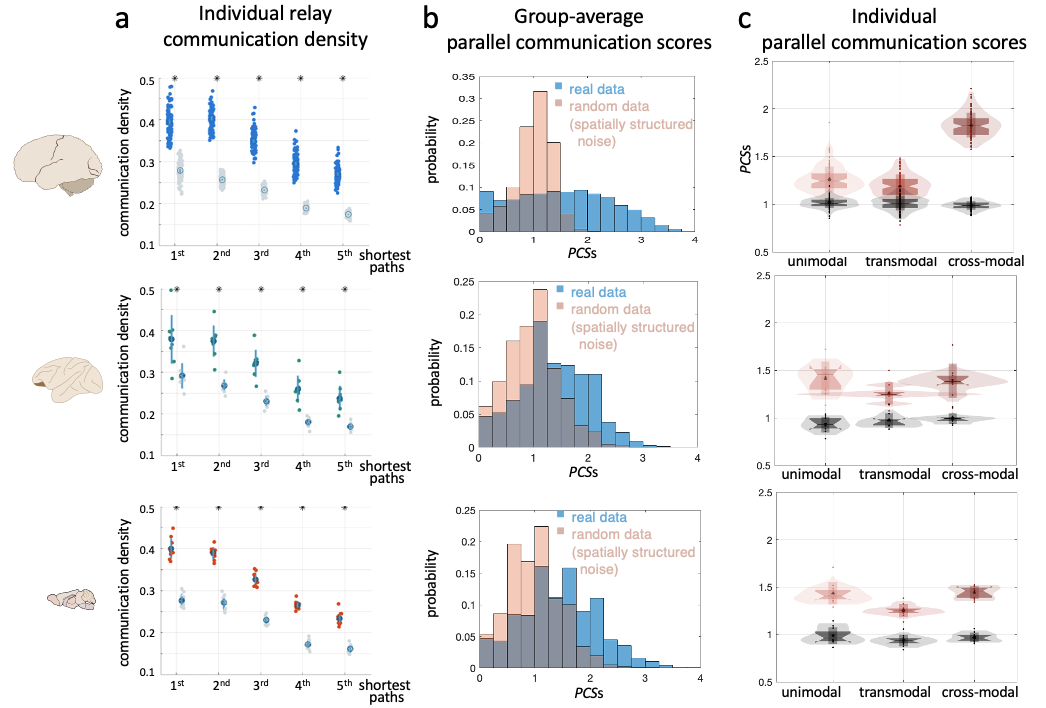

**Supplementary Figure 20. Comparison between real data and spatially structured physiological noise null model.** Left: schematic of human, macaque, and mouse brains from scidraw.io (https://doi.org/10.5281/zenodo.3925945, https://doi.org/10.5281/zenodo.3926117, https://doi.org/10.5281/zenodo.3925909). Each row in the figure corresponds to one species. (a) Box plots representing the percentage of short structural paths in individual brain networks respecting the data processing inequality (communication density). Each coloured dot represents an individual. Grey dots represent null distributions obtained by shuffling individual mutual information values while preserving network-level autocorrelation patterns (i.e., preserving the mutual information variogram); the brain network structural topology was unchanged (Methods). Circles and vertical bars indicate mean ± one standard deviation across individuals and randomizations. Paths are grouped according to the 1^st^ up to the 5^th^ shortest path between region pairs. (b) Histograms of group-average parallel communication scores (PCSs) obtained from real data (blue) and spatially structured noise (pink). For all the species, the real PCS histograms significantly differ from the null ones (two-sample Kolmogorov-Smirnov tests, p < .05), indicating that parallel communication levels are not the trivial by-product of the network-level autocorrelation present in functional brain data. (c) Average PCSs between unimodal systems, between transmodal systems, and between unimodal and transmodal systems (cross-modal communication) for individual experimental subjects. In the box plots, each dot represents a subject individual; vertical bars indicate mean ± standard deviation; notch bars indicate median and 1^st^-3^rd^ quartiles; shaded areas indicate 1^st^-99^th^ percentiles. Values from the spatially structured null model are reported in the same panels as grey box plots. For all species and brain systems, the real individual PCS significantly differ from the null ones, indicating that the network-level spatial autocorrelation of functional signals cannot explain system-dependent spatial communication patterns. Humans: n=100 biologically independent subjects; macaques: n=9 biologically independent subjects; mice: n=10 biologically independent subjects.

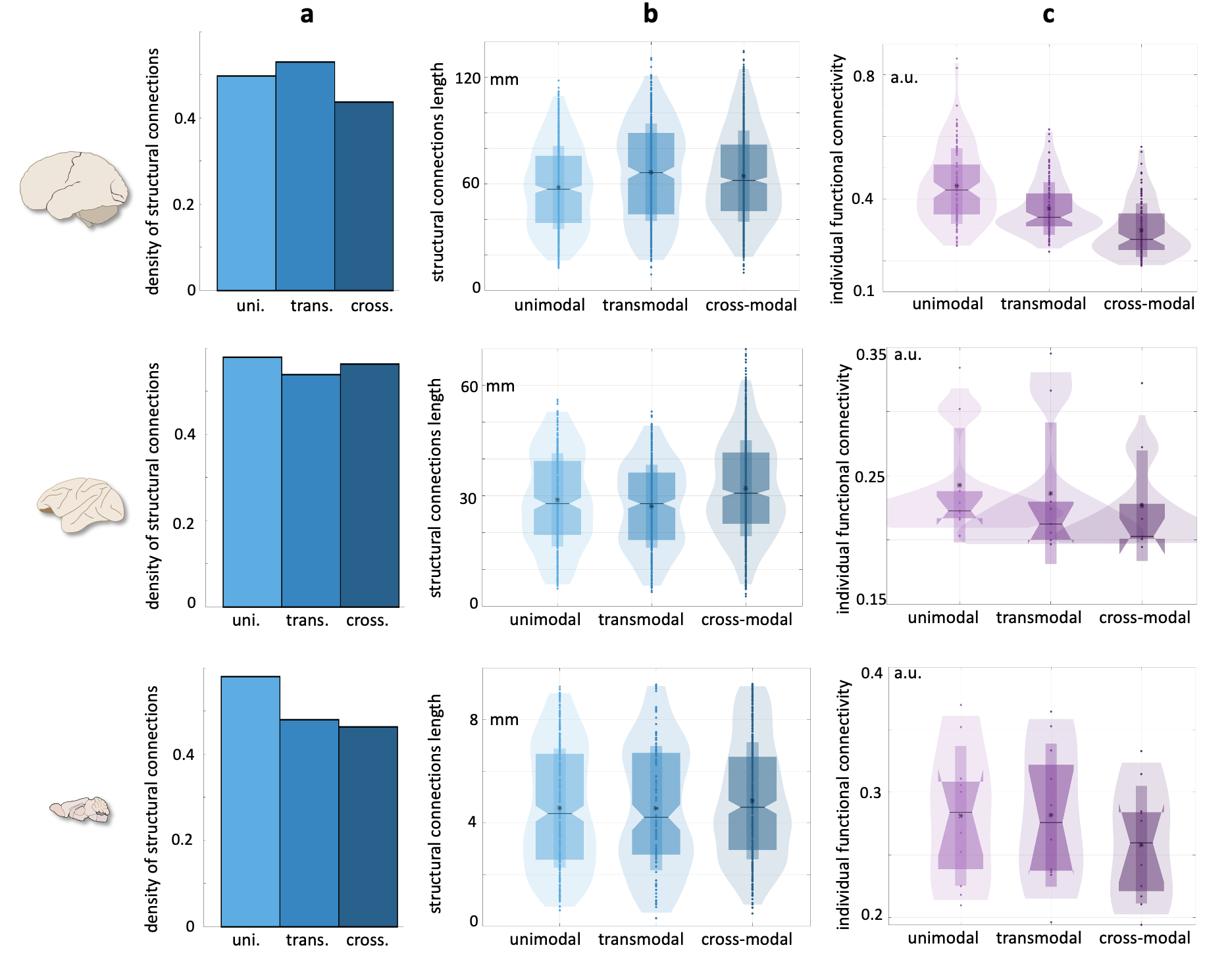

**Supplementary Figure 21. Structural and functional connectivity features across unimodal and transmodal systems.** Left: schematic of human, macaque, and mouse brains from scidraw.io (https://doi.org/10.5281/zenodo.3925945, https://doi.org/10.5281/zenodo.3926117, https://doi.org/10.5281/zenodo.3925909). Each row in the figure corresponds to one species. Density of structural connections (a), average length of structural connections (quantified as Euclidean distance between regions’ centroids in mm) (b), and average functional connectivity in individual subjects (quantified as mutual information between regions’ fMRI time series) (c) between unimodal systems, between transmodal systems, and between unimodal and transmodal systems (cross-modal communication). In (b) each dot represents a structural connection; in (c) each dot represents a subject. In the box plots, vertical bars indicate mean ± standard deviation; notch bars indicate median and 1^st^-3^rd^ quartiles; shaded areas indicate 1^st^-99^th^ percentiles. Cross-species characteristics of parallel communication patterns, with strong parallel communication streams between unimodal and transmodal (‘cross-modal’) areas in humans (represented in Fig. 3c), are not trivially explained by the number of anatomical connections, brain spatial embedding, and resting-state functional coupling alone. Humans: n=100 biologically independent subjects; macaques: n=9 biologically independent subjects; mice: n=10 biologically independent subjects. uni.=unimodal; trans.=transmodal; cross.=cross-modal; a.u.=arbitrary units.

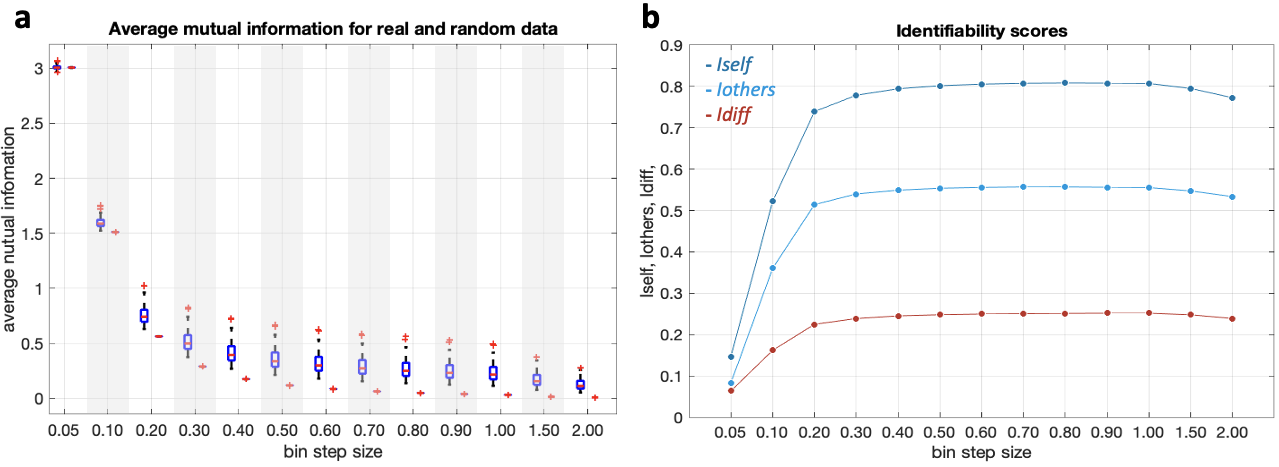

**Supplementary Figure 22. Choice of bin size for fMRI time series probability mass function estimation.** The bin size for the fMRI time series probability mass function estimation and mutual information computation was chosen by comparing real and null mutual information values, with null values obtained from multivariate gaussian data, and by assessing the fingerprinting accuracy of mutual information across bin sizes. (a) Brain-average mutual information values and average mutual information values for multivariate gaussian data with zero mean and diagonal covariance matrix. Mutual information values are reported in arbitrary units. For each bin step size, the boxplot on the left represents brain data of n=100 biologically unrelated human subjects; the boxplot on the right represents null data (100 null datasets). Note how the null boxplots are smaller than the real ones, denoting less data variability. Also notice how too small bin sizes can erroneously inflate mutual information values in null data. As expected, after a certain bin step size, the average mutual information values from real data are larger than the ones from null data, whereas the null data values go to zero. (b) Identifiability scores for different bin sizes: Iself (dark blue), Iothers (light blue), Idiff (red)^10^ (Methods), reported in arbitrary units. The fMRI time series of n=100 biologically unrelated human subjects were split into two parts of equal duration (400 time points each) and considered as test and retest data. Mutual information matrices were computed for test and retest data and the average within-subject (Iself) and between-subject (Iothers) test-retest Pearson’s correlation values were computed for each bin size. Idiff is the difference between Iself and Iothers: larger Idiff scores indicate better subject identifiability from mutual information data. We observe that for small bin sizes, the average mutual information values are high but the subject identifiability is low. Contrarily, for large bin sizes, the subject identifiability is high but the average mutual information values are low. We chose a bin size of 0.50 as a trade-off between relatively large mutual information values and near-maximal subject identifiability.

|  | Description | N | Anesthesia | MRI field | fMRI voxel size (mm) | fMRI TR (ms) | fMRI time points |
| --- | --- | --- | --- | --- | --- | --- | --- |
| h-HCP ^(a)^ | healthy human adults (46 males; 29.1 ± 3.7 years) | up to 100 | no | 3T | 1.6x1.6x1.6 | 720 | 800 (up to 1200) |
| q-NCS ^(b)^ | adult macaques (Macaca mulatta; 7 males; 8.4 ± 2.4 years) | 10 | no | 4.7T | 1.2x1.2x1.2 | 2600 | 500 |
| q-TVB ^(c)^ | male adult macaques (8 Macaca mulatta, 1 Macaca fascicularis) | 9 | yes, maintained with 1–1.5% isoflurane | 7T | 1.0x1.0x1.1 | 1000 | 600 |
| m-GG ^(d)^ | male adult mice (wild-type; 10 males; <6 months old) | 10 | no | 7T | 0.23x0.23x0.60 | 1000 | 600 (up to 1800) |
| m-AD3 ^(e)^ | male adult mice (wild-type; 10 males; 3 months) | 10 | yes, maintained with 0.5% isoflurane | 11.75T | 0.22x0.25x0.50 | 1000 | 600 |
| m-CSD1 ^(f))^ | male adult mice (wild-type; 51 males; 3 months) | up to 51 | yes, 3.5% isoflurane | 9.4T | 0.22x0.25x0.50 | 1000 | 360 |

**Supplementary Table 1. Cross-species datasets.**

N = number of participants

TR = repetition time

Sources:

^(a)^ Human Connectome Project (HCP), U100 dataset, HCP900 data release ^1^

^(b)^ INDI PRIMatE Data Exchange initiative, Newcastle University dataset (http://fcon_1000.projects.nitrc.org/indi/PRIME/newcastle.html) ^4^

^(c)^ The Virtual Brain ^3^

^(d)^ (https://data.mendeley.com/datasets/np2fx99hn6/2) ^9^

^(e)^ Mouse_rest_3xTG dataset (https://openneuro.org/datasets/ds001890/versions/1.0.1) ^5,6^

^(f)^ Grandjean et al., 2016 ^7^

|  | Beta | SE | t statistic | p-value |
| --- | --- | --- | --- | --- |
| Intercept | 2.27 | 0.26 | 8.71 | 4.7 ⋅ 10^-16^ |
| Species (macaque) | -3.40 | 0.35 | -9.75 | 3.7 ⋅ 10^-19^ |
| Species (mouse) | -4.31 | 0.59 | -7.24 | 5.9 ⋅ 10^-12^ |
| SC - number of connections (degree) | 0.57 | 0.06 | 0.11 | 3.3 ⋅ 10^-17^ |
| SC - average connection length | -0.68 | 0.19 | -3.53 | 4.9 ⋅ 10^-4^ |
| FC - MI sum | -1.14 | 0.17 | -8.01 | 4.7 ⋅ 10^-14^ |

**Supplementary Table 2. Relationship between parallel communication and connectome architecture.** Multiple regression models including group-average nodal parallel communication scores (PCSs, as depicted in Fig. 3a) as dependent variable, and the species (human, macaque and mouse encoded as two linearly independent regressors) and three connectome nodal features (the number of structural connections (structural degree); the average length of structural connections; the sum of functional connectivity weights) as independent variables (model *F*-statistic = 34.70, p < 10^-15^; *R*-squared = 0.42). All variables were z-scored. The length of structural connections was quantified as Euclidean distance between regions’ centroids (mm); functional connectivity weights were quantified as group-average mutual information between fMRI time series. SC=structural connectivity; FC=functional connectivity; MI=mutual information; Beta=estimated coefficient; SE=standard error of estimated coefficient; t=t statistic for Beta values; p-value=p-value from two-sided t test.

|  | Humans | | Macaques | | Mice | |
| --- | --- | --- | --- | --- | --- | --- |
| PCS Threshold | **SR*_low-PCS_*** | **SR*_high-PCS_*** | **SR*_low-PCS_*** | **SR*_high-PCS_*** | **SR*_low-PCS_*** | **SR*_high-PCS_*** |
| 1.20 | **72.0%** (0.9±1.0%) | **84.0%** (1.0±1.0%) | **44.4%** (10.1±9.4%) | **77.8%** (11.1±10.7%) | **40.0%** (10.1±9.3%) | **40.0%** (9.7±9.6%) |
| 1.30 | **73.0%** (0.9±1.0%) | **85.0%** (1.0±1.0%) | **66.7%** (10.7±10.5%) | **66.7%** (12.0±11.1%) | **40.0%**  (10.4±9.8%) | **40.0%** (9.9±9.6%) |
| 1.40 | **76.0%** (0.9±1.0%) | **81.0%** (1.0±1.0%) | **66.7%** (10.8±9.9%) | **77.8%** (10.6±10.9%) | **40.0%** (10.2±9.8%) | **40.0%** (10.6±9.9%) |
| 1.50 | **77.0%** (1.0±1.0%) | **79.0%** (0.9±1.0%) | **77.8%** (11.3±10.3%) | **77.8%** (10.9±10.7%) | **40.0%** (9.9±9.3%) | **30.0%** (9.1±8.9%) |
| Species median | **77.0%** (1.0±1.0%) | **80.0%** (1.0±9.5%) | **66.7%** (11.1±10.1%) | **77.8%** (11.4±10.7%) | **40.0%**  (10.1±9.7%) | **40.0%** (10.2±9.9%) |

**Supplementary Table 3. Individual identifiability from parallel communication scores: contribution of selective and parallel processing.** The table reports the participant identification success rate obtained when considering only the brain region pairs with, on average, a parallel communication score (PCS) lower than a threshold (SR*_low-PCS_*) or higher than a threshold (SR*_high-PCS_*). Success rates obtained from randomized data are reported in parenthesis (mean±standard deviation). Different PCS thresholds have been tested, including the median value of the group-average parallel communication score matrices of the three species. In the latter case, the threshold for the three species were the following: humans = 1.52; macaques = 1.33; mice = 1.40. Humans: n=100 biologically independent subjects; macaques: n=9 biologically independent subjects; mice: n=10 biologically independent subjects
